# Tactile Stimulation Improves Cognition, Motor, and Anxiety-Like Behaviours and Attenuates the AD Pathology in Adult APP ^*NL-G-F/NL-G-F*^ mice

**DOI:** 10.1101/2022.02.28.482218

**Authors:** Shakhawat R. Hossain, Hadil Karem, Zahra Jafari, Bryan E. Kolb, Majid H. Mohajerani

## Abstract

Alzheimer’s Disease (AD) is one of the largest health crises in the world. There are, however, limited but expensive pharmaceutical interventions to treat AD and most of the treatment options are not for cure or prevention, but to slow down the progression of the disease. The aim of this study was to examine the effect of tactile stimulation on AD-like symptoms and pathology in APP ^*NL-G-F/NL-G-F*^ mice, a mouse model of AD. The results show that tactile stimulation improves the AD-like symptoms on tests of cognition, motor, and anxiety-like behaviours and these improvements are associated with reduced AD pathology in APP mice.

## Introduction

Alzheimer Disease (AD) is a neurodegenerative brain disorder that causes cognitive and motor skills deficits. These behavioural symptoms are associated with the formation of extracellular Aβ plaques and intracellular tau phosphorylated proteins (Marcello et al., 2015), shrinkage of the cerebral cortex, hippocampus (Hpc), and basal ganglia (Pini et al., 2016), reduction of acetylcholine (Fischer et al., 1989), synaptic loss (Hamos et al., 1989), and disrupted gamma oscillations (Iaccarino et al., 2016) in the brain.

AD is difficult to treat and pharmacological treatments are not always effective in slowing the progression of the disease. Sensory stimulation, including tactile, auditory, visual, and olfactory, have been proposed as treatments for neurological disorders like AD. The benefits of these rehabilitation strategies are: 1) they are non-invasive; 2) they are cost effective; and, 3) they are easily translatable from preclinical studies to human clinical trials. For the purpose of this study, we focused on the beneficial effects of tactile stimulation (TS) in treating AD. Forms of TS ranging from skin-skin contact for new born infants, to gentle massage therapy for adults, have been proven to be beneficial for infant brain development and recovery from adult injury respectively (Gibb, Gonzalez, Wagenest, & Kolb, 2010; Kolb & Gibb, 2010). The receptors at the end of hair follicles and the dendrites in corpuscles in dermal and epidermal regions produce action potentials as a haptic response from TS. In addition, application of TS may also influence the peripheral nervous system (PNS), activating many endogenous mechanisms. Although the mechanism of TS in brain plasticity is not yet well understood, research shows that TS releases fibroblast growth factor-2 (FGF-2) (Gibb, 2005; Gibb et al., 2021), which crosses the blood brain barrier (BBB), and helps with neurogenesis, repair of nerve cells, cellular proliferation, survival, migration, and differentiation.

In this study, we aimed to assess cognitive, motor, and anxiety-like behaviours, and AD-like pathology, such as Aß plaques and hippocampal volume, to determine the effect of TS on adult APP^*NL-G-F/NL-G-F*^ mice, a mouse model of AD. Our prediction was that TS would enhance the cognitive, motor, and anxiety-like behaviours, and that these improvements would be associated with reduced Aß plaques and larger hippocampal volume.

## Methods and Materials

### Animals

Mice were housed in the Canadian Center for Behavioral Neuroscience (CCBN) vivarium, and all the behavioral, brain anatomical, and physiological tests and analyses were approved by the University of Lethbridge Animal Welfare Committee. APP^*NL-G-F/NL-G-F*^ (amyloid β-protein precursor), AD transgenic mice carrying Swedish (NL), Arctic (G), and Beyreuther/Iberian (F) mutations (Saito et al., 2014) provided by RIKEN Brain Science Institute were used in this research project. Nine females and six male APP^*NL-G-F/NL-G-F*^ (APP) adult transgenic mice, and six females and 6 male C57BL/6J (C57) were used in this project. All mice were given access to food and water ad libitum by the animal care staff. The mice were maintained on a 12-hour light and 12 hours dark cycle in a 21°C temperature controlled room in the vivarium. All training and behavioral testing was performed by the same experimenter during the light phase.

### Experimental Design

Mice from both APP and C57 strains were randomly assigned to four groups consisting of APP with tactile stimulation (APP-TS) group, APP without tactile stimulation (APP-NTS), C57BL/6J with tactile stimulation (C57-TS) group, and C57BL/6J without tactile stimulation (C57-NTS) group as per Figure1. Based on our earlier studies we did not expect to see a sex difference but we did include both sexes. Each group consists of a minimum 3 male and 3 female mice. There were no sex differences on any measure so we collapsed sex leaving n’s of 6 or more.

**Figure 1.**
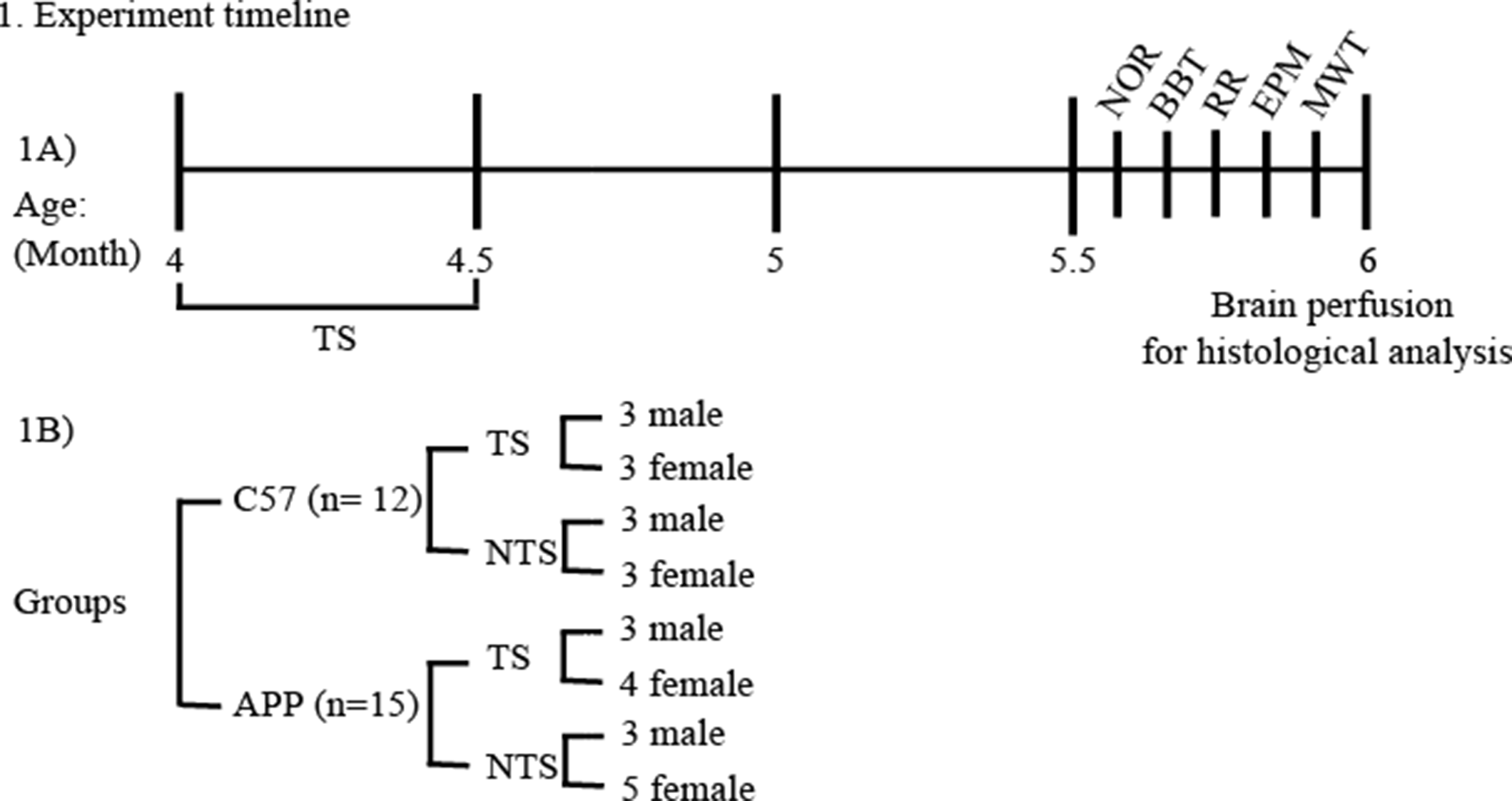
(A) Shows the experiment timeline in months (age of mouse). Animals were sacrificed a day after finishing the behavioral tests for Aβ quantifications. Aβ, amyloid-beta; NOR, novel object recognition; BBT, balance beam test; RR, rotarod test; EPM, elevated plus maze; MWT, Morris water task. (B) Shows the number of groups and number of male and female C57 and APP mice in each group.

### Tactile Stimulation Procedure

All the mice were handled for 5 minutes twice a day for 5 days prior to 4 months old. Only the TS groups of APP and C57 mice received manual TS at 4 months of age by lightly massaging each mouse with an experimenter’s fingers for 15 minutes with a frequency of 3 times a day for 15 days (8 am, 12 pm, and 4 pm). We applied TS when the APP mice were 4 months old, because the earliest onset of Aβ plaque formation in APP mice is ~3 months of age (Jafari et al., 2017; Mehla et al., 2019). At 6 months the Aβ plaque formation is completely saturated in the brain and deficits in cognition, motor and anxiety-like behaviour are also associated with the Aβ plaque formation. Therefore, we decided to apply TS at 4 months to determine if TS improves cognition, motor, and anxiety-like behaviours at 5.5 months and Aβ plaque formation at 6 months of age.

### Behavioral Tests

Several behavioral tests were performed at 5.5 months to measure the effect of TS on cognitive and motor functions. The balance beam (BB), rota-rod (RR), novel object recognition (NOR), activity box (AB), elevated plus maze (EPM), and the Morris water task (MWT) were conducted respectively by the same examiner with an alternating order of animals.

### Novel Object Recognition (NOR) Test

The NOR test was conducted to observe and measure the short term memory in the mice. Each mouse was placed in the same open field arena of 47cm x 50cm x 30cm with 2 similar objects for 5 minutes. After a 3-minute break, each mouse was exposed to one old and one novel object and the activity was recorded for 3 minutes. The time (seconds) spent with each old and novel object was manually recorded for analysis (Jafari et al., 2017). The discrimination index (DI) was calculated by using the formula (time spent with novel object-time spent with old object)/total time spent with both novel and old objects) (Ennaceur & Delacour, 1988).

### Morris Water Test (MWT)

The MWT task was performed to measure the spatial navigation abilities of the mice. Each mouse was placed in a 153cm diameter pool filled with water (23-25°C). The pool was located in a room with distal cues and virtually subdivided into 4 quadrants with starting points at north, west, east, and south. A hidden platform was placed in one fixed quadrant and was submerged ~1.0 cm during all 8 training days. Non-toxic white tempura paint was added to the pool water to make the water opaque, so that the mice would not have been able to see the platform. Each mouse was trained with 1 trial from each quadrant per day for 8 consecutive days (Water2100 Software vs.7, 2008). During each trial, the mouse was placed in the tank and each trial was stopped either once the mouse reached the platform, or if the mouse was unable to find the platform in 60 seconds. Data were recorded using an automated tracking system (HVS Image, Hampton, U.K.) and swim time (sec), swim speed (m/s), and swim distance (m) were calculated for analysis. On day 9, a probe trial was conducted, during which the platform was removed, and each mouse was allowed to swim freely for 60 seconds. For the analysis of probe trial, the time spent in the quadrant where the platform was located during training days was measured (Jafari et al., 2017).

### Balance Beam (BB) Test

The BB test was performed to measure the motor skills of each mouse. To conduct this test, the mice were trained to traverse across a 1 cm diameter, 100 cm long beam, which was 50 cm above a foam pad to cushion falling mice, to reach an escape box. On day 1, each mouse was trained for 3 successful trials. On day2, the traverse activity for each mouse was recorded for 3 trials and manually scored for the mean latency (sec), distance travelled (cm), and number of foot slips for analysis (Jafari et al., 2018; Tamura et al., 2012).

### Rotarod (RR) Test

The RR test was performed to measure the motor skills and the strength of gait in each mouse. All the mice were trained to walk on an automated 4 lane RR treadmill (ENV-575 M Mouse, Med Association Inc) on day 1. On day 2, each mouse was placed on the RR treadmill at 8rpm and 16rpm constant speed and at a 4-40 rpm alternating speed and recorded for 3 trials and the time (sec) each mouse was able to stay on the RR treadmill was recorded (Brooks and Dunnett, 2009).

### Elevated Plus Maze (EPM) Test

The EPM is a measure of anxiety-like behaviour in mice. The EPM apparatus was constructed from black Plexi-glass, which had two closed arms and two open arms. It was 40 cm high and two open arms were 5 cm wide and 27 cm long. The two closed arms were 10 cm wide, 40 cm long, and had 40 cm high walls. Each mouse was placed in the center of the EPM facing the closed arms. A camera was set up above the maze to film each mouse for 5 minutes. Each mouse was manually scored for time spent in the open arms (seconds), time spent in closed arms (seconds), number of entries to open arms, and number of entries to closed arms (Jafari et al., 2017). The EPM ratio was calculated by subtracting the number of entries to open arms from the number of entries to closed arms, divided by the total number of entries to both open and closed arms (Jafari et al., 2018).

### Quantification of Aβ plaque Area and Numbers

The methoxy-04 solution was prepared by diluting methoxy-X04 into 10% dimethyl sulfoxide, 45% propylene glycol, and 45% sodium phosphate saline. A 5mg/ml prepared methoxy-X04 was placed on a rotator at 4°C for 24 hours for better saturation, and the solution was stored at 4°C prior to the use. Methoxy-X04 was injected intraperitoneally at a dose of 10mg/kg using a 27 ½ G needle 24 hours before the perfusion of each animal (Bisht et al, 2016). Methoxy-X04, a fluorescent dye that selectively binds to β-pleated sheets found in Aβ plaques, has stronger specificity in staining Aβ plaques (Hefendehl et al. 2011).

The mice were perfused after the completion of the behavioral tests at the age of 6 months. Each mouse was injected with .05mg/kg of pentobarbital intraperitoneally. Then each brain received trans-cardial perfusion with 1x PBS until the blood ran clear followed with 4% PFA and the brain was extracted and post fixed with 4% PFA at 4°c for 24 hours. The brains were then transferred to 30% sucrose for solidification at least 48 hours before slicing with a cryostat machine with a thickness of 50μm. A Nanozomer fluorescent machine was used to colour the plaques and tangles in each brain section for analysis.

Each brain section was imaged automatically by using the Hamamatsu Nanozoomer 2.0-HT Scan System (Hamamatsu Photonics. Hamamatsu Japan) with a .23 μm/pixel resolution for quantification of Aβ plaques. The Ilastik 1.3.2rc2 and ImageJ 1.4.3.67 software were used for the plaque quantification. There were six coronal sections (Bregma: ~, +3.20, +2.96, +0.98, −2.06, −3.08, and −5.34 mm) that were selected corresponding to the mouse brain atlas (Paxinos and Franklin 2001) to quantify the total number of Aβ plaques and total plaque area (%) in each mouse brain (Saito et al. 2014). Five additional brain regions of interest (ROI’s): isocortex (IC), olfactory area (OA), medial-prefrontal cortex (mPFC), nucleus accumbens (NA), hippocampal area (HR) from each brain were selected for Aβ plaque quantifications (Jafari et al., 2017, 2018).

### Results and Statistical Analysis

All statistical analyses were performed using SPSS Statistics 24.0 at a significance level of 0.05 or better. None of the behavioural tests showed sex differences (P>05) so these data were collapsed across sex. Two way ANOVA was done for each behavioural test. The Bonferroni post-hoc test was used for each behavioural test, due to similar variance in each groups. The Bonferroni post-hoc analysis compares the means among multiple groups to determine significant differences between groups, while taking experimental errors into consideration. Results reported as mean ± S.E.M. Asterisks indicate *P<0.05 or **P<0.01 or ***P<0.001 value and partial eta squared (η2) indicates the effect size.

### NOR Test

The APP mice spent significantly less time with the novel object compared to all of the other groups. TS significantly improved the performance on novel object exploration in both C57 and APP mice. The overall significant ANOVA results comparing the 4 groups were: novel object time (F (3, 23) = 33.054, P≤ .0001, □2 = .812, power = 1.000), discrimination index ratio (F (3, 23) = 27.209, P≤ .0001, □2 = .780, power = 1.000).

The novel object time was significantly higher in C57-TS compared to C57 (F (1, 10) = 10.402, P≤ .012, □2= .565, power= .806) groups, in C57 compared to APP (F (1, 12) = 34.371, P≤ .0001, □2= .741, power= 1.000), and in APP-TS related to APP (F (1, 13) = 21.735, P≤ .0001, □2 = .750, power = .999) mice. The discrimination index ratio was higher in C57-TS mice relative to C57 (F (1, 10) = 7.722, P = .024, □2 = .491, power = .683), in C57 mice relative to APP (F (1, 12) = 25.974, P≤ .0001, □2 = .684, power = .997), and in APP-TS mice related to APP (F (1, 13) = 24.509, P≤ .0001, □2 = .690, power = .994). No significant difference was observed in time spent with the old object among the groups (F (3, 23) = .172, P = .914, □2= 0.22, power= .077) (Figure 2). A Bonferroni post-hoc analysis revealed that the C57-TS group spent significantly more time with novel object in comparison with C57 (P = .008), APP (P≤ .0001), and APP-TS (P≤ .0001) and the APP group spent significantly reduced amount of time with novel object in comparison with APP-TS (P = .001), C57 (P≤ .0001), and C57-TS (P≤ .0001). The highest discrimination index ratio was observed in the C57-TS in comparison with APP (P≤ .0001) and the lowest discrimination index ratio was observed in the APP group relative to APP-TS (P≤ .0001), C57 (P≤ .0001), and C57-TS (P≤ .0001) as per Bonferroni post hoc analysis.

**Figure 2.**
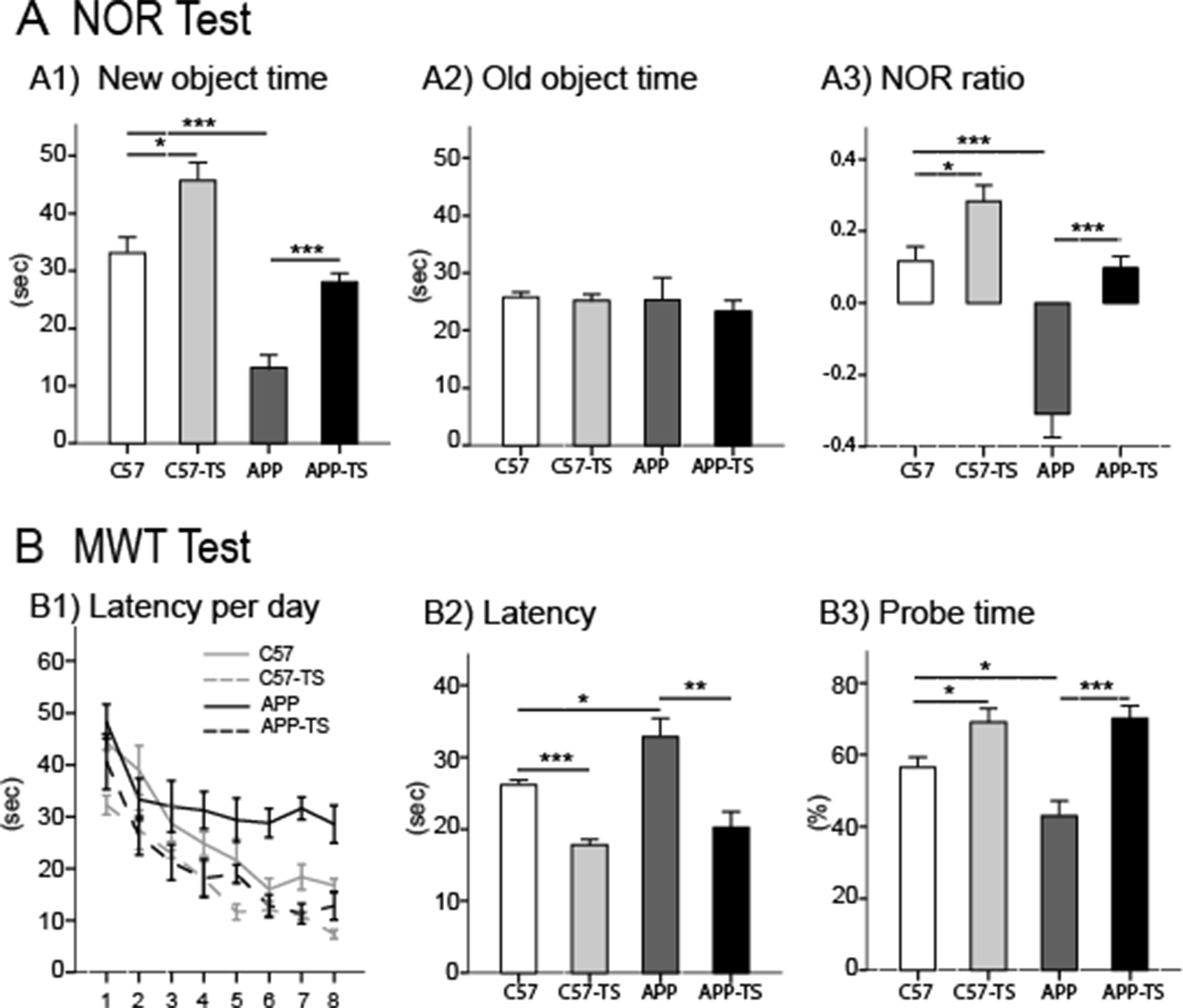
Results of cognitive tests. (A) NOR test: C57-TS showed a significantly longer new object time (sec) and higher NOR ratio, but no significant difference in old object time (sec) compared to C57, APP, APP-TS groups. (B) MWM test: The APP group showed significantly increased latency (sec) and shorter probe time (%) compared to C57, C57-TS, and APP-TS groups. Results reported as mean ± S.E.M. Asterisks indicate P*≤ .05, P**≤ .01, and P***≤ .001. NOR, novel object recognition; MWT, Morris water task.

**Figure 3.**
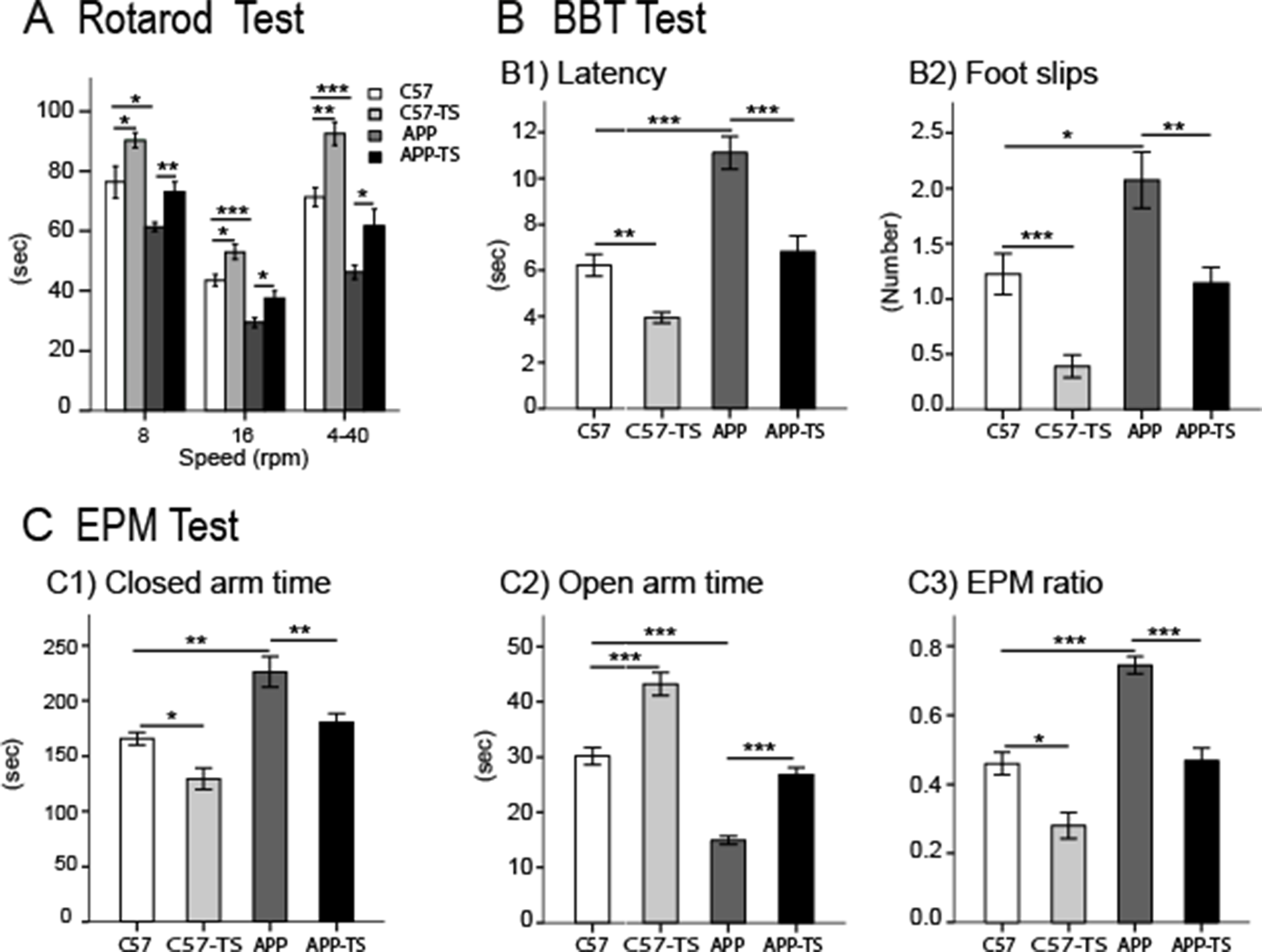
Results of motor tests (A and B) and anxiety-like behaviour test (C). (A) Rotarod test: A longer time spent on the rotarod (sec) with all three speeds in the C57-TS group compared to C57, APP, and APP-TS groups. (B) BBT: The APP group spent significantly longer time (sec) and higher number of foot slips compared to APP_TS, C57, and C57-TS groups. (C) EPM test: The APP group spent significantly increased amount of time (sec) in closed arms, decreased amount of time in open arms, increased EPM ratio for old object compared to APP-TS, C57, and C57-TS groups. Results reported as mean ± S.E.M. Asterisks indicate P*≤ .05, P**≤ .01, and P***≤ .001. BBT, balance beam test; EPM, elevated plus maze.

**Figure 4.**
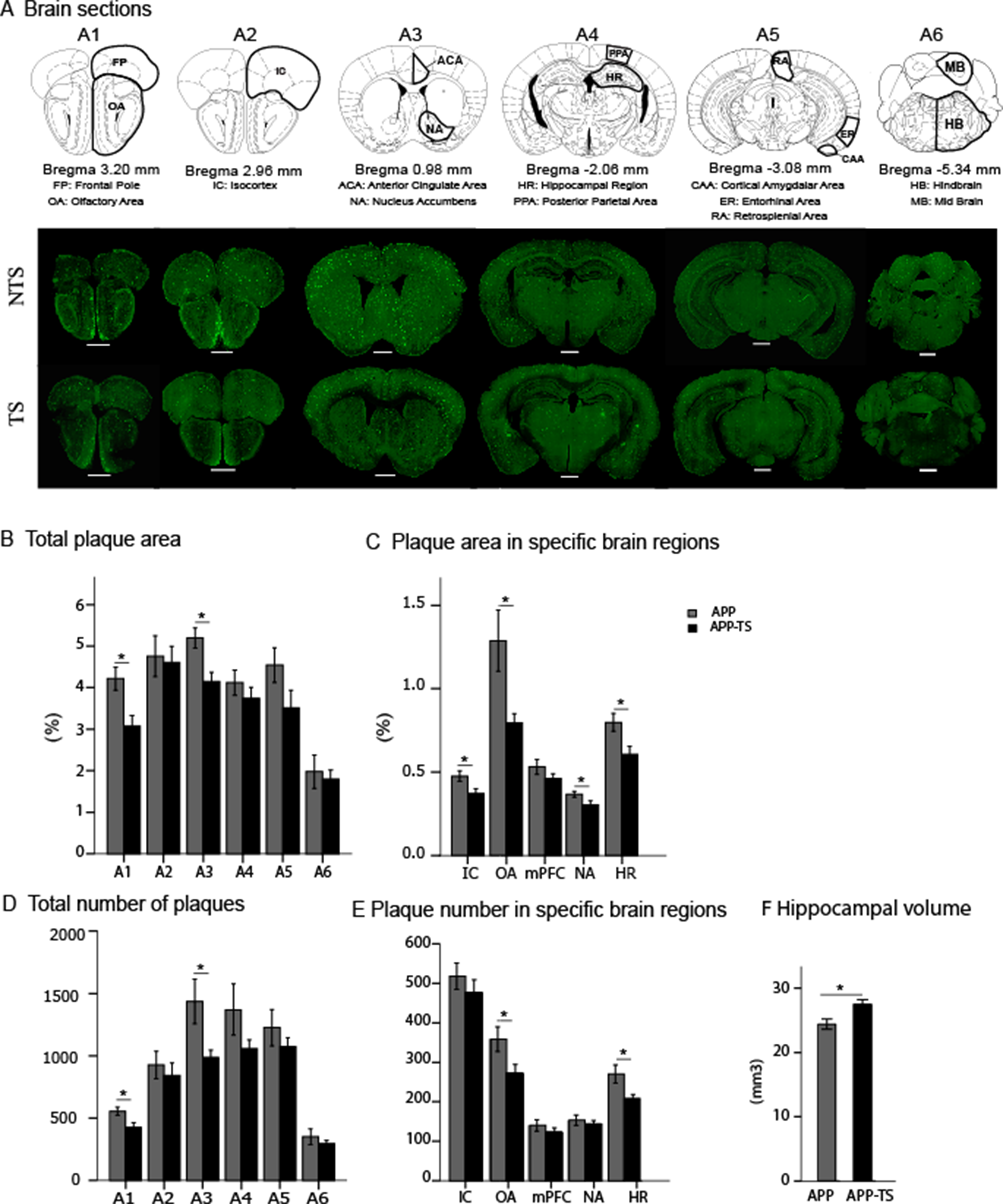
The Aβ plaque quantification in APP mice at the age of 6 months. (A) Six coronal brain sections (A1-A6: Bregma 3.20, 2.96, 0.98, −2.06, −3.08, and −5.34 mm) as a reference and Examples of experimental brain sections from both TS and NTS groups. (B) Total plaque area (%): The APP-TS group had significantly lower plaque area (%) in brain sections A1 and A3 compared to APP mice. (C) Plaque area (%) in specific brain regions: The APP-TS mice had significantly lower Plaque area (%) in IC, OA, NA, and HR compared to APP mice. (D) Total number of Plaques: The APP-TS group had significantly lower number of plaques in A1 and A3 brain sections compared to APP mice. (E) Plaque number in specific brain regions: The APP-TS group had significantly lower number of plaque number in OA and HR areas compared to the APP mice. (F) Hippocampal Volume: the hippocampal volume was significantly larger in the APP-TS group compared to APP mice. Aβ, amyloid beta; HR, hippocampal region; IC, isocortex; mPFC, medial prefrontal cortex; NA, nucleus accumbens; OA, olfactory area. Results reported as mean ±S.E.M. Asterisks indicate P*≤ .05, P**≤ .01, and P***≤ .001. Scale bar: 1 mm.

### MWT test

The APP mice were significantly slower to locate the sub-merged platform, showed a longer swim distance, and a reduced probe time in the target quadrant than each of the other groups. TS significantly improved the performances on all three measures for both of the C57 and APP mice.

The overall significant effects among all of the 4 groups are: latency (F (3, 23) = 12.377, P≤ .0001, □2 = .662, power = .998), swim distance (F (3, 23) = 24.008, P≤ .0001, □2 = .791, power = 1.000), and probe time (F (3, 23) = 12.385, P≤ .0001, □2 = .662, power = .998). During training days, swim latency was significantly decreased in the C57-TS mice compared to C57 (F (1, 10) = 56.858, P≤ .0001, □2 = .877, power = 1.000), in the C57 mice compared to APP (F (1, 12) = 4.859, P = .048, □2 = .288, power = .527), and in APP-TS mice compared to APP (F (1, 13) = 14.642, P = .003, □2 = .571, power = .935). The swim distance during the training days was also significantly decreased in the C57-TS mice relative to C57 (F (1, 10) = 19.843, P = .001, □2 = .665, power = .979), in the C57 mice relative to APP (F (1, 12) = 19.746, P = .001, □2 = .622, power = .982), and in the APP-TS relative to (F (1, 13) = 33.075, P≤ .0001, □2 = .750, power = .999). During the probe day, the amount of time spent in the target quadrant was significantly higher in the C57-TS mice compared to the C57 (F (1,10) = 5.737, P = .043, □2 = .418, power = .557), in the C57 compared to APP (F (1,12) = 6.087, P = .03, □2 = .337, power = .621), and in the APP_TS compared to the APP (F (1,13) = 25.741, P≤ .0001, □2 = .701, power = .996). No significant differences were observed in terms of the swimming speeds among the groups (F (3, 23) = .358, P = .784, □2 = .053, power = .107).

A Bonferroni post hoc analysis revealed that the C57-TS mice took significantly less time to locate the hidden platform during training days in comparison with the APP mice (P≤ .0001) and the APP mice took significantly more time than the C57-TS (P≤ .0001) and APP-TS (P = .001) mice. According to Bonferroni post hoc analysis the C57-TS mice swam the shortest distance during the training days compared to the APP (P≤ .0001) and the APP mice swam the longest distance compared to the C57 (P = .02), C57-TS (P≤ .0001), and APP-TS (P≤ .0001) mice. During the probe day, the C57-TS mice spent the highest amount of time in the target quadrant than the APP mice (P = .001) and the APP mice spent the least amount of time in the target quadrant than the C57-TS (P = .001), and the APP_TS (P≤ .0001) mice.

### BB Test

The APP mice were significantly slower to traverse the beam and made more slips than each of the other groups. TS significantly improved performance on both measures for both the C57 and APP mice.

The overall significant differences among all four groups in latency was F (3, 23) = 25.420, P≤ .0001, □2 = .761, power = 1.000, and number of foot slips is F (3, 23) = 12.398, P≤ .0001, □2 = .608, power = .999. The C57-TS group took significantly less time to cross the beam (F (1, 10) = 15.142, P = .005, □2 = .654, power = .924) and exhibited a reduced number of foot slips in C57 (F (1, 10) = 25.005, P = .001, □2 = .758, power = .991) compared to C57 group. The C57 mice also took significantly less time to cross the beam (F (1, 10) = 25.857, P≤ .0001, □2 = .665, power = .997) and had a reduced number of foot slips F (1, 12) = 5.996, P = .029, □2 = .316, power = .620) in comparison to APP. In contrast, the APP group took significantly longer to cross the beam (F (1, 13) = 18.133, P = .001, □2 = .564, power = .977) and showed an increased number of foot slips (F (1, 13) = 8.744, P = .01, □2 = .384, power = .785) compared to APP-TS. A Bonferroni post-hoc analysis revealed that the APP group took the longest time to cross the beam relative to C57 (P≤ .0001), C57-TS (P≤ .0001), and APP-TS (P≤ .0001) and had the highest number of foot slips relative to C57 (P = .05), C57-TS (P≤ .0001), and APP-TS (P = .019). The C57-TS group took significantly shorter time to traverse the beam relative to APP (P≤ .0001) and APP-TS (P = .044), and had significantly reduced number of foot slips compared to APP (P≤ .0001) as per Bonferroni post hoc analysis.

### RR Test

The APP group showed significantly impaired performance compared to APP-TS, C57, and C57-TS groups and C57-TS group exhibited the most improved performances in all RR speeds (8 rpm, 16 rpm, and 4-40 rpm) among the groups (Figure 5). TS significantly improved the RR performances in both C57-TS and APP-TS groups in relative to C57 and APP respectively.

The overall significant differences between all four groups were at 8 rpm: F (3, 23) = 13.779, P≤ .0001, □2 = .646, power = 1.000; at 16 rpm: F (3, 23) = 21.735, P≤ .0001, □2 = .739, power = 1.000; and at 4-40 rpm: F (3, 23) = 25.446, P≤ .0001, □2 = .768, power = 1.000. Compared to C57 group, C57-TS mice showed significantly improved performance in all three RR speeds, i.e., 8 rpm: F (1, 10) = 5.661, P = .039, □2 = .361, power = .575; 16 rpm: F (1, 13) = 8.421, P = .016, □2 = .457, power = .744; and 4-40 rpm: F (1, 10) = 18.442, P = .002, □2 = .648, power = .971. Similarly, APP-TS group exhibited significantly improved performance in all three RR speeds, i.e., 8 rpm: F (1, 13) = 12.029, P = .004, □2 = .481, power = .893; 16 rpm: F (1, 13) = 8.155, P = .014, □2 = .385, power = .752; and 4-40 rpm: F (1, 13) = 7.388, P = .018, □2 = .362, power = .710 compared to APP mice. In contrast, the APP group exhibited impaired performances in all three RR speeds, i.e., 8 rpm: F (1, 12) = 8.865, P = .013, □2 = .446, power = .773; 16 rpm: F (1, 12) = 36.463, P≤ .0001, □2 = .768, power = 1.000; and 4-40rpm: F (1, 12) = 41.175, P≤ .0001, □2 = .789, power = 1.000 in relative to C57 mice. A Bonferroni post-hoc analysis revealed that the C57-TS group showed significantly improved performances compared to all other groups at all three speeds, i.e., 8 rpm: APP (P = .035), and APP-TS (P = .016); 16 rpm: APP (P≤ .0001), and APP-TS (P = .001), and 4-40 rpm: C57 (P = .016), APP (P≤ .0001), and APP-TS (P≤ .0001).

### EPM Test

The C57-TS mice were significantly less anxious, and spent significantly more time in the open arms of the maze and less time in the closed arms of the maze compared to the other experimental groups. TS reduced the anxiety like behavior in both C57 and APP mice. The overall significant effects noted between all of the four groups were: open arm time (F (3, 23) = 67.143, P≤ .0001, □2 = .914, power = 1.000), closed arm time (F (3, 23) = 16.092, P≤ .0001, □2 = .718, power = 1.000), and EPM ratio (F (3, 23) = 31.905, P≤ .0001, □2 = .834, power = 1.000).

The open arm time was significantly higher in C57-TS mice compared to C57 (F (1, 10) = 23.049, P = .001, □2 = .742, power = .986), in C57 compared to APP (F (1, 12) = 88.735, P≤ .0001, □2 = .881, power = 1.000), and in APP-TS compared to APP (F (1, 13) = 74.454, P≤ .0001, □2 = .871, power = 1.000). In contrast, the closed arm time was significantly lower in C57-TS mice compared to C57 (F (1, 10) = 10.516, P = .012, □2 = .568, power = .810), in C57 compared to APP (F (1, 12) = 12.492, P = .004, □2 = .510, power = .900), and in APP-TS compared to APP (F (1, 13) = 9.675, P = .01, □2 = .468, power = .808). In addition, the EPM ratio was significantly lower in C57-TS mice relative to C57 (F (1, 10) = 10.534, P = .012, □2 = .568, power = .811), in C57 related to APP (F (1, 12) = 50.867, .0001, □2 = .809, power = 1.000), and APP-TS related to APP (F (1, 13) = 41.357, .0001, □2 = .790, power = 1.000). A Bonferroni post-hoc analysis revealed that the C57-TS group had the longest open arms time in comparison with C57 (P≤ .0001), APP (P≤ .0001), and APP-TS (P≤ .0001) and the APP group had the shortest open arms time in comparison with APP-TS (P≤ .0001), C57 (P≤ .0001), and C57-TS (P≤ .0001). In contrast, the C57-TS spent the lowest time in the closed arm related to APP (P≤ .0001) and APP-TS (P = .023), and APP spent the highest time in the closed arms relative to C57 (P = .003), C57-TS (P≤ .0001), and APP-TS (P = .023) as per Bonferroni post-hoc analysis. A Bonferroni post hoc analysis for EMP ratio discovered that the C57-TS showed lowest EMP ratio for closed arms compared to C57 (P = .014), APP (P≤ .0001), and APP-TS (P = .007) and highest EMP ratio for closed arms in the APP mice relative to C57 (P≤ .0001), C57-TS (P≤ .0001), and APP-TS (P≤ .0001).

### Impact of TS on the amyloid-β (Aβ) plaque pathology

The deposition of total number of Aβ plaques was higher in all 6 coronal sections and TS attenuated the formation of Aβ plaques in the APP mice. Although the pattern of increased amount of Aβ deposition was observed in all 6 coronal positions of the APP mice compared to APP-TS, it was significantly higher in sections + 3.20 (F (1,10) = 5.885, P = .041, □2 = .424, power = .568), and + 0.98 (F (1,10)= 6.529, P =.034, □2 = .449, power = .612). In addition, there was a trend for the total number of Aβ plaques to be higher in APP mice compared to APP-TS (P = .073).

TS also positively influenced the formation of Aβ by reducing the area of plaques (%) in APP mice. Again, the reduced pattern of the area of Aβ plaques (%) in all 6 coronal positions were observed; however, the area of Aβ plaques (%) was significantly smaller in + 3.20 (F (1, 10) = 7.729, P = .024, □2 = .491, power = .684), and + 0.98 (F (1, 10) = 8.455, P = .02, □2= .514, power = .722). In addition, the total area of Aβ plaques (%) was significantly reduced in APP-TS mice compared to APP (F (1, 10) = 9.991, P = .013, □2 = .555, power = .790).

The positive influence of TS on Aβ plaque area (%) was also observed in ROI’s. The reduced pattern of Aβ plaques area (%) was observed in all the ROI’s but CAA and HB of APP-TS group compared to APP mice. However, APP-TS mice showed significantly reduced Aβ plaque area (%) in IC (F (1, 10) = 6.148, P = .038, □2 = .435, power = .586), OA (F (1, 10) = 6.183, P =.038, □2 = .436, power = .589), and HR (F (1, 10) = 7.321, P = .027, □2 = .478, power = .660), compared to APP group. Furthermore, the number of Aβ plaques was significantly reduced in APP-TS group in OA (F (1, 10) = 5.044, P = .049, □2 = .335, power= .527) and HR (F (1, 10) = 5.884, P = .036, □2 = .370, power = .591).

## Discussion

There are three main findings from this investigation: 1) TS ameliorated the cognitive and motor dysfunctions and reduced anxiety-like behavior; 2) TS attenuated the Aβ plaque size and numbers; and 3) TS enlarged the hippocampal volume in adult APP mice. We consider each finding in turn.

### The impact of TS on cognition and motor learning and anxiety-like behavior

#### Cognition

Impaired learning and memory is one of the common symptoms of AD in humans, and our findings from this study as well as the previous studies conducted in our lab (Karem, 2019; Jafari et al., 2018; 2019) demonstrate a similar impairment in APP mice. The goal of this study was to establish the influence of TS in reducing the symptoms and pathology of AD in APP mice. Our findings from both the MWT and NOR tests suggest that TS improves cognition not only in APP mice, but also in C57 mice, which is the wild-type of APP mice. In the MWT test, both C57 and APP mice that received TS displayed significantly shorter latency and distance, and longer probe time, suggesting improvement in their spatial learning and memory as seen in previous studies using TS to stimulate recovery in brain-injured animals (Angeles et al., 2016; Kolb & Gibb, 2010). Similarly, there was a significant increase in time spent with the novel object and less time with old object in the NOR test showing that both groups that received TS demonstrated enhanced short-term memory. TS has been proven to be beneficial to treat depression-like symptoms in rats as TS positively influences the HPA axis (Angeles et al., 2016), increases the level of neurotrophic factors such as BDNF in the hippocampus, increases GFAP signaling (Antoniazzi et, al., 2016 and Roversi et al., 2019), prevents hippocampal damage due to neonatal hypoxia in rats (Rodrigues et al. 2004), and increases secretion of acetylcholine (ACh) in the hippocampus of rats (Dudar et al., 1979). TS in the form of maternal licking and grooming increases the brain-derived neurotrophic factor (BDNF) mRNA, NMDA receptors, improved spatial learning and memory in rats (Liu et al., 2000).

#### Motor Skills

A deterioration in motor skills is a common symptom of AD in humans. The findings from the current study also show similar motor deficits as previously shown in studies on APP mice (Jafari et al., 2018; 2019). The results from both BB and RR test revealed that TS significantly improved the performances in both motor tests. In BB test both the C57 and APP mice that received TS traversed the balance beam faster, and had fewer foot slips, which indicates improved balance and motor coordination. Likewise, both groups that received TS showed markedly improved performances on the rotating wheel, suggesting enhancement of their motor coordination as well. Studies of TS on rats with medial prefrontal cortical lesions have previously shown improvements in a skilled reaching task (Kolb & Gibb, 2010; Gibb et al., 2010). Similarly, the application of TS has been proven to be beneficial in improving motor recovery in human stroke victims (Hunter et al., 2008) and motor development in preterm infants (Field et al., 1986). Numerous studies have shown that TS increased response to somatosensory stimulation in the sensory motor cortex (Schaechter, 2011), dendritic length in frontal and sensorimotor cortex (Gibb et al., 2010), recovery of 20 Hz rebound in motor-cortical excitability (Parkkonen et al., 2018), and sensorimotor rhythm-based brain-computer interface performance (Shu et al., 2018). TS has also been shown to be beneficial in improving locomotion and exploratory behavior, as well as reducing protein carbonyl levels in the cortex, hippocampus, and sub-thalamic regions (Boufleur, et al., 2012). Application of gentle message therapy also increased urine dopamine by 31% (Field et al., 2009). These changes are important because enhanced neuro-synaptic plasticity in frontal and sensorimotor cortex, dopamine, and motor-cortical excitability plays very vital role in motor balance and coordination.

#### Anxiety-like behavior

Anxiety-like behavior, due to stress and depression, has been identified as a risk factor for AD (Aznar and Knudsen, 2011). Anxiety may lead to frustration and possibly continue throughout the progress of AD. In this study, we aimed to determine the positive effect of TS on anxiety-like behaviour in APP mice. Our findings from the EPM test indicated that TS significantly reduced the anxiety in both C57 and APP mice, as these mice spent more time in the open arms and had an increased EPM ratio. Studies on rodents have shown that TS reduces anxiety-like behavior (Freitas et al., 2015; Boufleur, et al., 2012), increases the responsiveness to drugs such as benzodiazepine (Boufleur, et al., 2012), and reduces the sensitization of psychostimulant drugs such as amphetamine (Mouhammad et al., 2010). Studies on either prenatal or postnatal TS have been shown to alter cortical thickness and striatum size (Muhammad and Kolb, 2011), increase plasma antioxidant compounds such as vitamin C and glutathione peroxidase in the cortex, hippocampus, and sub-thalamic region (Boufleur, et al., 2012), and lower plasma cortisol level (Jafari et al, 2018; 2019). Field et al. (2009) reviewed the studies on the positive impacts of massage therapy on humans and concluded that massage therapy reduced saliva cortisol by 31%, and increased urine serotonin by 28%. Reduced cortisol and increased serotonin play a very essential role in improving anxiety-like behaviour.

### The impact of TS on Aβ pathology in APP adult mice

The loss of cholinergic neurons, atrophy of hippocampal regions, the neocortex, and thalamus, and formation of tau-proteins, tangles, and are a few of the neural symptoms of AD. In this study we investigated the effect of TS on Aβ pathology, and the hippocampal volume in APP mice. Although the formation of Aβ plaques was significantly reduced in some brain regions, but not all, a reduction pattern of Aβ plaques was observed throughout the brains that received TS. One of the very first senses that begins to diminish is olfaction in early stages of AD patients (Kovács et al., 2001) and similar finding have been established in APP mice as well (Jafari et al., 2018). In our findings, the biggest significant anatomical difference observed was the reduced number and size of Aβ plaques in OA of the mice that received TS (Figure 6A). The formation of Aβ plaques is also visible in most parts of the neocortex, and hippocampal regions (Jafari et al., 2018) of APP mice. In this study, we demonstrated that TS significantly reduced Aβ plaque numbers and size in the hippocampus and isocortex.

We also observed a significant reduction of the percentage of Aβ plaque areas and Aβ plaque numbers in bregma position + 3.2 mm and + 0.98 mm, and a pattern of decreased Aβ plaque numbers and in the percentage of Aβ plaque areas shown in all other coronal positions of the mouse of brains in the mice that received TS. A collapse across all the coronal planes revealed a significant reduction of the percentage of Aβ plaque areas in the mice that received TS. Further analysis of ROI’s revealed a significant reduction of the percentage of Aβ plaque areas. A recent research by Martorell et al., (2019) shows that auditory and visual stimulation reduce Aβ plaque in the neocortex and hippocampus and improve spatial and recognition memory in 5XFAD mice.

### The impact of TS on hippocampal volume (Hpc) in APP adult mice

Research on humans (Gosche et a., 2002) and rodents (Zahra et al., 2017 and 2018) has shown that one of the main hallmarks of AD is the shrinkage of hippocampal volume. We were able to show that application of TS in early stages of AD, prevents the hippocampal volume from shrinking in APP mice. Similarly, along with the larger hippocampal volume, there was a reduced Aβ plaque number, and reduced percentage of Aβ plaque area, which was associated with improved cognitive and motor skills in APP mice that received TS.

## Conclusion

Although TS has been successfully implemented in various clinical settings ranging from premature infants, institutionalized infants, work places, wound care, and treating HIV, this study the first to use this intervention in APP mice to counter the progression of AD pathology. Our findings demonstrate that TS improves cognitive and motor functions and anxiety-like behaviour in APP mice and these improved functions are associated with reduced Aβ plaque areas and numbers and increased hippocampal volume in their brain. These results suggest that TS, which is a non-invasive and cost-effective intervention, could be applied to human AD patients, even after symptoms are obvious. These findings offer promise for the application of TS in patients with AD. However, further research is required to discover the brain mechanisms regarding changes in the gene expression, electrophysiology, neurotransmitters, FGF-2, and synapses in response to TS in both neurologically normal and APP mice.

## Acknowledgements

This work was supported by Natural Sciences and Engineering Research Council of Canada (NSERC) Discovery Grant #40352 to MHM, Alberta Innovates (MHM), Alberta Alzheimer Research Program (MHM), Alzheimer Society of Canada (MHM), Alberta Prion Research Institute (MHM), Canadian Institute for Health Research (MHM), and Alberta Registered Nurse Education Trust (SRH). We thank Dr. Takashi Saito and Prof. Takaomi C Saido from “Laboratory for Proteolytic Neuroscience RIKEN Center for Brain Science, Wako-shi, Saitama, Japan” for providing the APP^NL-G-F/NL-G-F^ mice as a gift. We also thank Di Shao for animal breeding.

Writers would like to acknowledge the grant supported by

## Authors Contribution

S.H., Z.J., M.H.M., and B.E.K. designed and conceptualized the study. M.H.M., and B.E.K. supervised the study. S.R.H. performed the behavioural experiments. S.R.H. analyzed the behavioural data. S.R.H., and H.K. performed the immunohistochemistry. H.K., analyzed the immunohistochemistry data. S.R.H., and B.E.K. wrote the manuscript. S.H., Z.J., M.H.M., and B.E.K. all commented on and edited the manuscript.

## References

Angeles, G.A.M., Carmen, R.O.M., Wendy, P.M., & Socorro, R. M. (2015). Tactile stimulation effects on hippocampal neurogenesis and spatial learning and memory in prenatally stressed rats. Brain Research Bulletin, Volume 124, Pages 1–11, https://doi.org/10.1016/j.brainresbull.2016.03.003

Antoniazzi, C. T. D., Metz, V. G., Roversi, K., Freitas, D. L., Vey, L. T., Dias, V. T., Segat, H. J., Duarte, M. M. M. F., & Burger, M. E. (2016). Tactile stimulation during different developmental periods modifies hippocampal BDNF and GR, affecting memory and behavior in adult rats. Hippocampus, Volume 27, Issue 2, https://doi.org/10.1002/hipo.22686

Aznar, S. & Knudsen, G. M. (2011). Depression and Alzheimer’s Disease: Is Stress the Initiating Factor in a Common Neuropathological Cascade? Journal of Alzheimer’s Disease, Volume 23, no. 2, 177–193, DOI: 10.3233/JAD-2010-100390

Bisht, K., El Hajj, H., Savage, J. C., Sanchez, M. G., & Tremblay, M. E. (2016). Correlative light and electron microscopy to study microglial interactions with beta-amyloid plaques. J Vis Exp 112.

Boufleur, N., Antoniazzi, C.T.D., Pase, C.S., Benvegnu, M.D., Barcelos, R.C.S., Dolci, G.S., Dias, V.T., Roversi, K., Koakoskira, G., Rosa, J.G., Barcellos, L.J.G., & Burger, M.E. (2012). Neonatal tactile stimulation changes anxiety-like behavior and improves responsiveness of rats to diazepam. Brain Research, Volume 1474, Pages 50–59, https://doi.org/10.1016/j.brainres.2012.08.002

Brooks, S. P., & Dunnett, S. B. (2009). Tests to assess motor phenotype in mice: a user’s guide. Nature Rev Neurosci. 10:519–529.

Bye, C.M., & McDolnald, R.J. (2019). A Specific Role of Hippocampal NMDA Receptors and Arc Protein in Rapid Encoding of Novel Environmental Representations and a More General Long-Term Consolidation Function. Front. Behav. Neurosci, https://doi.org/10.3389/fnbeh.2019.00008

Collingridge, G. (1987). The role of NMDA receptors in learning and memory. Nature News and Views, Volume 330, pages 604–605.

Comeau, W. Hastings, E. & B. Kolb, B. (2007). Pre-and postnatal fgf-2 both facilitate recovery and alter cortical morphology following early medial prefrontal cortical injury. Behav Brain Res, 180,pp. 18–27. https://doi:10.1016/j.bbr.2007.02.026

Dementia Statistics: Alzheimer’s Disease International. (2019). Retrieved September, 2019 from https://www.alz.co.uk/research/statistics

Dudar, J.D., Whishaw, I.Q., & Szerb, J.C. (1979). Release of acetylcholine from the hippocampus of freely moving rats during sensory stimulation and running. Neuropharmacology Volume 18, Issues 8-9, August-September 1979, Pages 673–678. https://doi.org/10.1016/0028-3908(79)90034-0

Ennaceur, A., & Delacour, J. (1988). A new one-trial test for neurobiological studies of memory in rats. 1: Behavioral data. Behav Brain Res, 31(1):47–59. https://doi.org/10.1016/0166-4328(88)90157-X

Field, T., Hernandez-Reif, M., Diego, M., Schanberg, S., & Kuhn, C. (2009). Cortisol decreases and serotonin and dopamine increases following massage therapy. International Journal of Neuroscience, Volume 115, Issue 10, 1397–1413. https://doi.org/10.1080/00207450590956459

Field, T.M., Schanberg, S.M., Scafidi, F., Bauer, C.R., Vega-Lahr, N., Garcia, R., Nystrom, J., & Kuhn, C.M. (1986). Tactile/Kinesthetic Stimulation Effects on Preterm Neonates. Pediatric, Volume77, issue 5.

Fisher, W., Gage, F. H., & Björklund, A. (1989). Degenerative Changes in Forebrain Cholinergic Nuclei Correlate with Cognitive Impairments in Aged Rats. European Journal of Neuroscience, Volume1, Issue1, pp. 34–35, https://doi.org/10.1111/j.1460-9568.1989.tb00772.x

Freitas, D., Antoniazzi, C.T.D., Segat, H.J., Metz, V.G., Vet, L.T., Barcelos, R.C.S., Duarte, T., Duarte, M.M.M.F., & Burger, M.E. (2015). Neonatal tactile stimulation decreases depression-like and anxiety-like behaviors and potentiates sertraline action in young rats. International Journal of Developmental Neuroscience, Volume 47, Part B, Pages 192–197, https://doi.org/10.1016/j.ijdevneu.2015.09.010

Gibb, R.L. (2005) This would be her thesis – I do not have it with me

Gibb, R.L., Gonzalez, C.L.R., Wegenast, W., & Kolb, B.E. (2010). Tactile stimulation promotes motor recovery following cortical injury in adult rats. Behavioural Brain Research, Volume 214, Issue 1, Pages 102–107, https://doi.org/10.1016/j.bbr.2010.04.008

Gibb, R.L., Kovalchuk, A., & Kolb, B. (2020). Tactile stimulation of functional recovery after perinatal cortical injury is mediated by FGF-2. In submission.

Gosche, K. M., Mortimer, J. A., Smith, C. D., Markesdery, W. R., & Snowdon, D. A. (2002). Hippocampal volume as an index of Alzheimer neuropathology Findings from the Nun Study. Neurology, 58 (10), DOI: https://doi.org/10.1212/WNL.58.10.1476

Hamos, J. E., DeGennaro, L. J., & Drachman, D. A. (1989). Synaptic loss in Alzheimer’s disease and other dementias. Neurology, 39 (3), DOI: https://doi.org/10.1212/WNL.39.3.355

Hefendehl, J. K., Wegenast-Braun, B. M., Liebig, C., Eicke, D., Milford, D., Calhoun, M. E., Kohsaka, S., Eichner, M., & Jucker, M. (2011). Long-term in vivo imaging of beta-amyloid plaque appearance and growth in a mouse model of cerebral beta-amyloidosis. J Neurosci. 31:624–629, DOI: https://doi.org/10.1523/JNEUROSCI.5147-10.2011

Herreros, F. (2013). An investigation of the attachment formation and organization of infants living in Chilean institutions. Published PhD thesis, The New School, ProQuest Dissertations Publishing, 2013. 3566444. https://search.proquest.com/docview/1418273384?accountid=12063

Hisselmo, M. (2006). The role of acetylcholine in learning and memory. Current Opinion Neurobiol, 16(6):710–5. http://doi:10.1016/j.conb.2006.09.002

Hunter, S.M., Crome, P., Sim, J., & Pomeroy, V.M. (2008). Archives of Physical Medicine and Rehabilitation, Volume 89, Issue 10, Pages 2003–2010, https://doi.org/10.1016/j.apmr.2008.03.016

Iaccarino, H. F., Singer, A. C., Martorell, A. J., Rudenko, A. Gao, F., Gillingham, T. Z., Mathys, H., Seo, J. Kritskiy, O., Abdurrob, F., Adaikkan, C., Canter, R. G., Rueda, R., Brown, E. N., Boyden, E. S., & Tsai, L. H. (2016). Gamma frequency entrainment attenuates amyloid load and modifies microglia. Nature, 540(7632):230–235, https://doi.org/10.1038/nature20587

Jafari, Z., Okuma, M., Karem, H., Mehla, J., Kolb, B., & Mohajerani, M. (2019). Prenatal noise stress aggravates cognitive decline and the onset and progression of beta amyloid pathology in a mouse model of Alzheimer’s disease. Neurobiology of Aging, 77, 66–86, https://doi.org/10.1016/j.neurobiolaging.2019.01.019

Jafari, Z., Mehla, J., Kolb, B., & Mohajerani, M. (2018). Gestational Stress Augments Postpartum ß-Amyloid Pathology and Cognitive Decline in a Mouse Model of Alzheimer’s Disease. Cerebral Cortex, Volume 29, Issue 9, Pages 3712–3724, https://doi.org/10.1093/cercor/bhy251

Kol, A., Adamsky, A., Groysman, M., Kreisel, T., London, M., & Goshen, I. (2019). Astrocytes Contribute to Remote Memory Formation by Modulating Hippocampal-Cortical Communication During Learning. BioRxiv, doi: https://doi.org/10.1101/682344

Kolb, B. & Gibb, R. (2011). Brain Plasticity and Behaviour in the Developing Brain. J Can Acad Child Adolesc Psychiatry, 20(4): 265–276, PMCID: PMC3222570 PMID: 22114608

Kolb, B. & Gibb, R. (2010). Tactile stimulation after frontal or parietal cortical injury in infant rats facilitates functional recovery and produces synaptic changes in adjacent cortex. Behavioural Brain Research, Volume 214, Issue 1, Pages 115–120, https://doi.org/10.1016/j.bbr.2010.04.024

Kovács, T.C.A., Cairns, N.J., & Lantos, P.L. (2001). Olfactory centres in Alzheimer’s disease: olfactory bulb is involved in early Braak’s stages. NeuroReport, Volume 12 - Issue 2 - p 285–288,

Lange, K. W., Guo, J., Kanaya, S., Lange, K. M., Nakamura, Y., & Li, S. (2019). Medical foods in Alzheimer’s disease. Food Science and Human Wellness, Volume 8, Issue 1, Pages 1–7, https://doi.org/10.1016/j.fshw.2019.02.002

Liu, D., Diorio, J., Day, J.C., Francis, D.D., & Meaney, M.J. (2000). Maternal care, hippocampal synaptogenesis and cognitive development in rats. Nature Neuroscience volume 3, pages799–806.

Marcello E, Gardoni F, Di Luca M. (2015). Alzheimer’s disease and modern lifestyle: what is the role of stress? J Neurochem. 134:795–798. https://doi.org/10.1111/jnc.13210

Martorell, A. J., Paulson, A. L., Suk, H., Abdurrob, F., Drummond, G. T., Guan, W., Young, Z. J., Kim, D. N., Kritskiy, O., Barker, S. J., Mangena, V., Prince, S. M., Brown, E. N., Chung, K., Boyden, E. S., Singer, A. C., & Tsai, L. (2019). Multi-sensory Gamma Stimulation Ameliorates Alzheimer’s-Associated Pathology and Improves Cognition. Cell, Volume 177, Issue 2, Pages 256–271.e22, https://doi.org/10.1016/j.cell.2019.02.014

Maruyama, K., Shimoju, R., Ohkubo, M., Maruyama, H., & Kurosawa, M. (2012). Tactile skin stimulation increases dopamine release in the nucleus accumbens in rats. The Journal of Physiological Sciences, Volume 62, Issue 3, pp 259–266, https://doi.org/10.1007/s12576-012-0205-z

Mehla, J., Lacoursiere, S. G., Lapointe, V., McNaughtin, B. L., Sutherland, R. J., McDonald, R. J., & Mohajerani, M. H. (2019). Age-dependent behavioral and biochemical characterization of single APP knock-in mouse (APPNL-G-F/NL-G-F) model of Alzheimer’s disease. Neurobiology of Aging, Volume 75, Pages 25–37, https://doi.org/10.1016/j.neurobiolaging.2018.10.026

Miranda, M., Kent, B.A., Morici, J. F., Gallo, F., Saksida, L.M., Bussey, T.J., Weisstaub, N., & Bekinschtein, P. (2018). NMDA receptors and BDNF are necessary for discrimination of overlapping spatial and non-spatial memories in perirhinal cortex and hippocampus. Neurobiology of Learning and Memory, Volume 155, November 2018, Pages 337–343, https://doi.org/10.1016/j.nlm.2018.08.019

Monfils, H., Driscoll, I., Melvin, N., & Kolb, B. (2006). Differential expression of basic fibroblast growth factor-2 in the developing brain. Neuroscience, 141, pp. 213–221. http://doi:10.1016/j.neuroscience.2006.03.047

Muhammad, A., Hossain, S., Pellis, S.M., & Kolb, B. (2011). Tactile stimulation during development attenuates amphetamine sensitization and structurally reorganizes prefrontal cortex and striatum in a sex-dependent manner. Behav Neurosci, 125(2): 161–74, http://doi:10.1037/a0022628

Parkkonen, E., Laaksonen, K., Parkkonen, L., & Forss, N. (2018). Recovery of the 20 Hz Rebound to Tactile and Proprioceptive Stimulation after. Neural Plasticity, Volume 2018, Article ID 7395798, 11 pages, https://doi.org/10.1155/2018/7395798

Paxinos G, and Franklin, K. B. J. 2001. The Mouse Brain in Stereotaxic Coordinates. San Diego: CA: Academic Press.

Pini, L., Pievani, M., Bocchetta, M., Altomare, D., Bosco, P., Cavedo, E., Galluzzi, S., Marizzoni, M., & Frisoni, G. B. (2016). Brain atrophy in Alzheimer’s Disease and aging. Ageing Research Reviews, Volume 30, Pages 25–48, https://doi.org/10.1016/j.arr.2016.01.002

Richards, S., Mychasiuk, R., Kolb, B., & Gibb, R. (2012). Tactile stimulation during development alters behaviour and neuroanatomical organization of normal rats. Behavioural Brain Research, Volume 231, Issue 1, Pages 86–91. https://doi.org/10.1016/j.bbr.2012.02.043

Ridley, R.M., Bowes, P.M., Baker, H.F., & Crow, T.J. (1984). An involvement of acetylcholine in object discrimination learning and memory in the marmoset. Neuropsychologia, Volume 22, Issue 3, 1984, Pages 253–263, https://doi.org/10.1016/0028-3932(84)90073-3

Rodrigues, A. L., Arteni, N.S., Abel, C., Zylbersztejn, D., Chazan, R., Viola, G., Xavier, L., Achaval, M., & Netto, C.A. (2004). Tactile stimulation and maternal separation prevent hippocampal damage in rats submitted to neonatal hypoxia-ischemia. Brain Research, Volume 1002, Issues 1-2, 26 March 2004, Pages 94–99, https://doi.org/10.1016/j.brainres.2003.12.020

Saito, T., Matsuba, Y., Mihira, N., Takano, J., Nilsson, P., Itohara, S., Iwata, N., & Saido, T. C. Single App knock-in mouse models of Alzheimer’s disease, Nat. Neurosci., 17 (2014), pp. 661–663, doi: 10.1038/nn.3697

Santello, M., Toni, N., & Andrea Volterra, A. (2019). Astrocyte function from information processing to cognition and cognitive impairment. Nature Neuroscience, volume 22, pages154–166, http://doi.org/10.1038/s41593.018.0325.8

Chaechter, J.D., Van Oers, C.A.M.M., & Groisser, B.N. (2011). Increase in Sensorimotor Cortex Response to Somatosensory Stimulation Over Subacute Poststroke Period Correlates With Motor Recovery in Hemiparetic Patients. Neurorehabilitation and Neural Repair, 26(4) 325–334, https://doi.org/10.1177/1545968311421613

Sheard, J. M. (2014). Malnutrition and Neurodegenerative Diseases. Curr Nutr Rep, 3: 102. https://doi.org/10.1007/s13668-014-0078-2

Shu, X., Chen, S., Meng, J.,Yao,L., Sheng, X., Jia, J., Farina, D., & Zhu, X. (2018). Tactile Stimulation Improves Sensorimotor Rhythm-Based BCI Performance in Stroke Patients. IEEE Transactions on Biomedical Engineering, Volume: 66, Issue: 7, 1987 – 1995, http://doi:10.1109/TBME.2018.2882075

Tamura, H., Ohgami, N., Yajima, I., Iida, M., Ohgami, K., Fujii, N., Itabe, H., Kusudo, T., Yamashita, H., & Kato, M. (2012). Chronic Exposure to Low Frequency Noise at Moderate Levels Causes Impaired Balance in Mice. PLOS ONE 7(6): e39807. https://doi.org/10.1371/journal.pone.0039807

